# An application of the MR-Horse method to reduce selection bias in genome-wide association studies of disease progression

**DOI:** 10.1101/2024.07.19.604143

**Authors:** Killian Donovan, Jason Torres, Doreen Zhu, William G Herrington, Natalie Staplin

## Abstract

Genome-wide association studies (GWAS) of disease progression are vulnerable to collider bias caused by selection of participants with disease at study entry. This bias introduces spurious associations between disease progression and genetic variants that are truly only associated with disease incidence. Methods of statistical adjustment to reduce this bias have been published, but rely on assumptions regarding the genetic correlation of disease incidence and disease progression which are likely to be violated in many human diseases. MR-Horse is a recently published Bayesian method to estimate the parameters of a general model of genetic pleiotropy in the setting of Mendelian Randomisation. We adapted this method to provide bias-reduced GWAS estimates of associations with disease progression, robust to the genetic correlation of disease incidence and disease progression and robust to the presence of pleiotropic variants with effects on both incidence and progression. We applied this adapted method to simulated GWAS of disease incidence and progression with pleiotropic variants and varying degrees of genetic correlation. When significant genetic correlation was present, the MR-Horse method produced less biased estimates than unadjusted analyses or analyses adjusted using other existing methods. Type 1 error rates with the MR-Horse method were consistently below the nominal 5% level, at the expense of a modest reduction in power. We then applied this method to summary statistics from the CKDGen consortium GWAS of kidney function decline. MR-Horse attenuated the effects of variants with known likely biased effects in the CKDGen GWAS, whilst preserving effects at loci with likely true effects.

## Introduction

Genome-wide association studies (GWAS) of disease progression are increasingly common due to the availability of sufficiently large cohorts of genotyped participants with specific diseases and long-term follow-up^1–3^. Selection for disease cases implicitly conditions on the processes causing incident disease, thus inducing biased associations through any confounders of the incidence - progression relationship^4,5^ (Fig 1). This bias may inflate type 1 error rates in GWAS of disease progression. Eliminating this bias by study design is challenging, as in order to define “disease progression”, participants must be stratified in some way by disease status. Measurement and elimination of this bias is important for understanding the genomic architecture of disease traits, and also could be applied to reduce the bias of estimates in non-genetic epidemiological studies.

**Figure 1:**
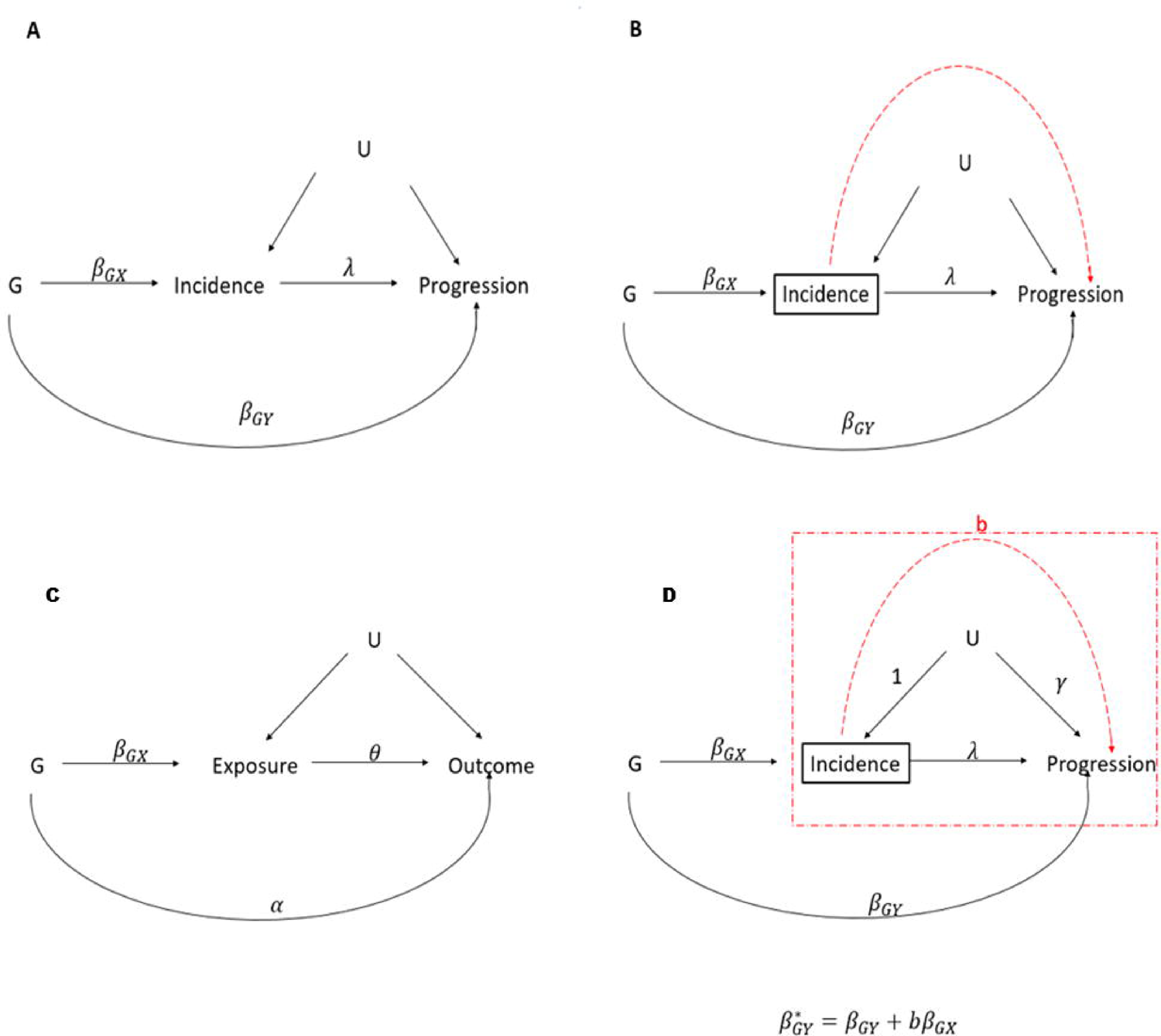
Conditioning on incident disease leads to biased associations of genetic variants with disease progression (A-B). Comparison of MR Horse model and its adaptation for mitigating collider bias (C – D). 1A: Directed acyclic graph (DAG) representing genetic effects on disease progression and incidence in the presence of confounders. 1B: Conditioning on incident disease (black rectangle) in GWAS of progression can induce biased associations through the confounders U. (red dashed line). G: participant genotype β_GX_: effect of genetic variant on disease incidence *β_GX_:* direct effect of genetic variant on disease progression, *U*: confounders of the incidence/progression relationship, λ: direct effect of disease incidence on disease progression 1C: DAG representing Mendelian Randomisation with pleiotropic variants, incorporating the notation used in original MR-Horse publication: G: participant genotype, *β_Gx_*: effect of genetic variant on disease exposure, α: direct (pleiotropic) effect of genetic variant on outcome, U: confounders of the incidence/progression relationship, θ: direct causal effect of exposure on outcome 1D: Adaptation of the DAG in 1A to illustrate the applicability of this model to the setting of selection bias in GWAS. λ: direct causal effect of disease incidence on disease progression, γ: effect of confounders on disease progression b: biasing pathway through U and γ, due to conditioning on Incidence (rectangle). Effects from *U* to Incidence are set as 1, allowing the relative confounder effects to vary by varying a single parameter γ.

**Figure 2:**
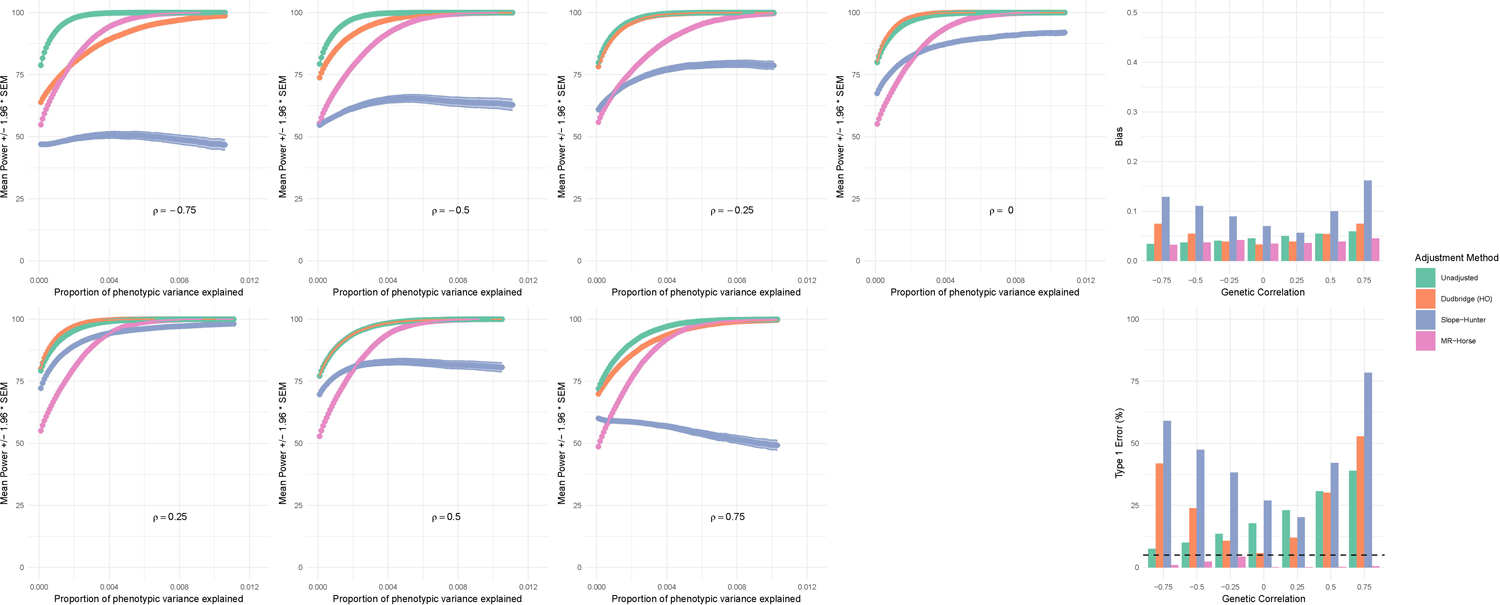
The adapted MR-Horse method has lower bias and type 1 error rates across the range of genetic correlations, at the expense of a reduction in statistical power. Left: Power of unadjusted and adjusted analyses to detect β_GY_ effects across the range of effect sizes, by different values of ρ. Plotted values are mean power (%) across 1,000 simulation iterations with shaded areas +/- 1.96 * SEM. Phenotypic variance explained per variant defined as: 2β^2^ *maf*(1 - *maf*). Right: Bias (top panel) and type 1 error rates (bottom panel) of each method (mean bias of *G_I_* and *G_IP_* variants [variants with direct effects on incidence] across 1,000 simulation iterations). Unadjusted analyses have increasing type 1 error rates with increasingly positive correlation. Dudbridge and Slope-Hunter analyses have increasing bias and type 1 error at extremes of correlation. MR- Horse retains control of type 1 error across range of correlation.

If individual-level genetic data are available, and the selection-process responsible for the bias is well understood, then the bias can be addressed by inverse-probability weighting of participants according to the probability of selection^6^. Unfortunately these data are often not available. Methods have been proposed to adjust GWAS summary statistics in order to measure and reduce selection bias^6–8^. Given that the bias arises from conditioning on disease incidence, these methods use knowledge of genetic variant associations with disease incidence to estimate the bias of each SNP estimate in the stratified progression GWAS.

The method proposed by Dudbridge and colleagues^7^ (termed “instrument effect regression” by some authors^8^) assumes that there is no correlation between direct genetic effects of a variant on incident disease (β*_GX_*) and its effects on disease progression (β*_GY_*), and thus any correlation between estimated effects on incidence and progression must be due to collider bias according to the equation

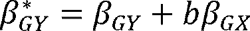

where β^*^*_GY_* denotes the biased value of β*_GY_* observed in a GWAS stratified by incident disease. The collider bias effect is thus defined by the coefficient *b*, which can be estimated by linear regression of estimated progression effects from the biased GWAS on estimated incidence effects from a separate study (provided incident disease in both studies is defined in an measurement error in β*_GX_*. The underlying assumption of no correlation is very strong, and in identical manner). They also propose two possible mechanisms for adjusting results for our view likely to be violated in most disease contexts. In the case of chronic kidney disease for example, incident disease is generally defined based on the measurement of a threshold value of a biomarker-based (usually serum creatinine) estimate of glomerular filtration rate (eGFR), and disease progression is typically defined as rate of change of this value over time (eGFR slope)^9^. In such an example, given that both phenotypes are defined based on the same measurements, it is likely that there is significant correlation between genetic effects on the phenotypes.

The method proposed by Mahmoud and colleague^10^ (Slope-Hunter method) does not assume that β*_GX_* and β*_GY_* are uncorrelated for all SNPs. Gaussian mixture models are used to cluster incidence-associated SNPs into a “hunted” cluster of SNPs with similar values of 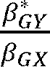 and a second cluster without such a relationship (the “pleiotropic” cluster). Any correlation between β*_GX_* and β*_GY_* among the “hunted” SNPs is assumed to be caused by the collider bias effect, which can thus be estimated by linear regression of estimated progression effects on estimated incidence effects in this cluster. Accurate identification of the appropriate “hunted” cluster requires that a plurality of variants with similar values of 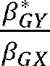 only have true effects on disease incidence and not disease progression. This is termed the **Ze**ro **M**odal **R**esidual **A**ssumption [ZeMRA], and is less restrictive than that of the Dudbridge method, but still likely to be violated in certain human diseases. It is therefore desirable to have a method which can reduce the bias of disease progression associations without assumptions regarding the underlying genetic correlation of the traits of interest. Given the analogies between this assumption of no correlation and the InSIDE (**In**strument **S**trength **I**ndependent of **D**irect **E**ffects) assumption of Mendelian Randomisation methods, we sought to apply an MR method robust to violations of the InSIDE assumption to the problem of selection bias in GWAS.

The MR Horse method developed by Grant and Burgess^11^ can give unbiased causal effect estimates in the Mendelian Randomisation setting, even in the presence of correlated shrinkage prior (horseshoe prior) on pleiotropic effects (α), to a set of summary statistics for pleiotropic effects (Fig 1C). This method uses a Bayesian approach to fit a model employing a effects to escape shrinkage, while shrinking most of the effects to 0. The assumptions variants associated with an exposure. This horseshoe prior allows variants with true pleiotropic effects and causal effect θ are constant across genetic variants. This model is very similar to the underlying this model are that effects in Fig 1C are approximately linear, and that confounder proposed model of causation in GWAS of disease progression (Fig 1D). The application of this model to disease progression GWAS can be summarized as follows, amending some of the notation used by Grant and Burgess:

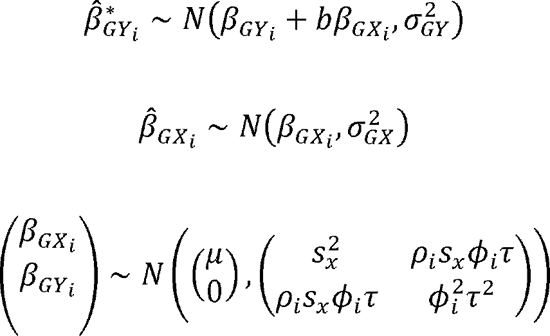

where ρ_i_ represents the correlation coefficient between β_GXi_ and β_GYi_ and Φ_i_ and are half Cauchy distributed variables representing the local and global shrinkage parameters respectively of a horseshoe prior^12^ for C. Applying this model to a set of summary statistics estimation of SNP effects on disease progression, robust to correlation between β_GY_ and β_GX_. for SNP associations with disease incidence and disease progression may allow unbiased collider bias vs in the MR setting. Firstly, in the original MR-Horse setting (Fig 1) represents There are some important differences in the interpretation of parameters in the setting of the direct causal effect of the exposure on the outcome. In the collider bias setting this value *b*, incorporates residual direct effects from “Incidence” to “Progression” not completely pathway through the confounders *U*. It therefore defines the magnitude of the collider bias but accounted for by adjustment for Incidence, in addition to the effects mediated along the is not intuitively interpretable in the same way as in the original MR-Horse formulation.

Secondly, in the MR setting α_i_ represents the pleiotropic effects of the genetic variant *i*, and its value is not of significant interest *per se*, whereas in the application to collider bias, β_GYi_ is the direct genetic effect of a variant on disease progression and thus is the estimand of interest. Finally, in the MR setting causal effects cannot be estimated when there is an effect of a SNP on a confounder *U*. This restriction also applies to the setting of disease progression GWAS. If a SNP has an effect on a confounder *U*, this effect will be captured in β_GX_ estimates and would exhibit a different correlation structure with β_GY_ than the direct β_GX_ effects, invalidating the MR-Horse model.

We aimed to adapt the MR-Horse method to produce bias-reduced estimates of disease-progression associations in GWAS which adjust for incident disease, robust to correlation between β_GX_ and β_GY_. We then tested this method in published GWAS of CKD progression, taking advantage of the fact that there are known biased associations of genetic variants with progression^13^ which can be used as negative controls.

## Methods

### Simulations

In order to test this novel application of the method we first simulated GWAS of disease incidence and progression, using knowledge of chronic kidney disease to approximately inform modelling choices. Genotypes were simulated for 10,000 individuals at 2,000 independent SNPs disease incidence or progression (α), 1% of SNPs had effects on disease incidence only (α), 1% in Hardy-Weinberg equilibrium^14^ and in linkage equilibrium. 95% of SNPs had no effect on of SNPs had effects on disease progression only (α) and 3% of SNPs had effects on both incidence and progression (α), with correlated β_GXi_ and β_GYi_ with correlation coefficient ρ (constant across *G_IP_* variants). This set-up produces a large pleiotropic cluster of variants explaining more of the phenotypic variance of disease progression than the non-pleiotropic cluster. This situation is likely to violate the assumptions of both the Dudbridge and Slope-Hunter methods, but is a plausible architecture of chronic kidney disease related traits. In order to approximate realistic distributions of effect sizes across the range of allele frequencies, minor allele frequencies were first drawn from a uniform distribution between 0.01 and 0.49, and standardised effect sizes were defined as 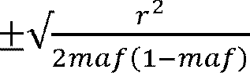 where variant level *r*^2^ values were drawn from the positive half of a normal distribution with mean 0. Confounders *r^2^*, were simulated as a random normal variable, and disease incidence (α) was initially simulated under a liability threshold model where the liability had a 40% SNP heritability according to the equation:

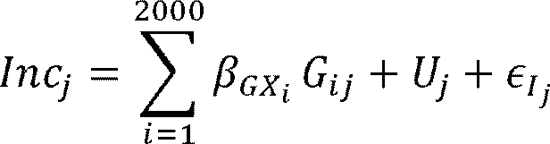

A threshold value of the phenotype was selected to define the binary phenotype of incident CKD such that the prevalence of CKD in the simulated population was 40%. Kidney function decline or disease progression was then simulated in all participants as a continuous variable with genetic effects on progression scaled such that the phenotype has 30-40% SNP heritability (estimated in GWAS among the entire simulated population) and 30-40% of its variance explained by the confounders *U*, according to the equation

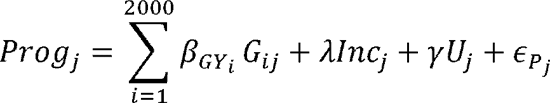

The values of λ and γ (Fig 1) were set as 0.1 and 1.5 respectively for all simulations. GWAS of the binary CKD incidence phenotype was then performed in the whole population (logistic simulated participants with incident CKD. Simulations were repeated with differing values of ρ.

We then applied Dudbridge’s method, the Slope-Hunter method, and the MR-Horse method to these GWAS results. For both Dudbridge and Slope-Hunter methods, only variants associated with disease incidence (we employed a threshold of *p* > 0.001) were used to estimate *b*. The Hedges-Olkin approximation was used to adjust Dudbridge estimates for measurement error, as the alternative simulation extrapolation methods performed unpredictably at extremes of genetic correlation. Confidence intervals for the Slope Hunter estimates were bootstrapped by knowledge of chronic kidney disease. The prior probability for ρ was 2*r* – 1 where *r ∽ Beta(3,1)* over 200 iterations. The prior probabilities for the MR-Horse model parameters were informed. This prior puts most of the probability weighting around high positive prior probability distribution for the global shrinkage parameter τ was set as *τ ∽ Cauchy+ (0,1)*, correlation but allows for some lower and even negative correlation for some variants. The prior probability distribution for the global shrinkage parameter and the prior for the local shrinkage prior φ_i_ was *τ ∽ Cauchy+ (0,1)*. Uninformative priors were used for *b ∽ Uniform+ (−10,10), μ ∽ N(0,1)*. Posterior parameter estimates were drawn from four parallel Markov Chains simulated in JAGS using 15,000 iterations with a 20% burn-in. Power, type 1 error and bias were recorded across 1,000 simulation iterations. For better comparison of method performance, power was the identification of a significant association of a *G_0_* or *G_1_* SNP with disease progression. measured across the entire range of simulated variant effect sizes. Type 1 error was defined as

Significant effects were defined at *p* < 0.05 for the regression based methods, and if the 95% equal-tailed credible interval for the MR-Horse estimate of β_GYi_ did not include 0. Bias was defined as |β_GYi_ - β_GYi_| for each method.

Sensitivity analyses were performed including simulated populations of different sizes, different relative cluster sizes (testing relative performances of the adjustment methods when assumptions of the other methods are met), and sparser or less sparse MR-Horse prior distributions for β*_GYi_* (Supplemental Data).

All analyses were performed in R 4.3.1 and JAGS 4.3.2 on the Oxford University Medical Sciences Division Biomedical Research Computing Cluster.

### Application to CKDGen data

To illustrate the performance of our method in a real dataset we used data from large meta-analyses of GWAS of kidney disease traits performed by the CKDGen consortium. We used summary statistics from two GWAS: (i) a GWAS of creatinine-based eGFR from Wuttke et al.^15^ and (ii) a GWAS of eGFR decline over time, adjusted for baseline eGFR from Gorski et al.^13^. As pointed out by the authors, adjustment for eGFR in GWAS (ii) should introduce a collider bias effect and spurious associations between genetic variants associated with cross-sectional eGFR alone and eGFR decline. Notably this scenario of adjustment for a baseline variable is different to our simulated scenario of stratification by incident disease, but the use of creatinine-based eGFR measurements as the phenotype of interest has the illustrative advantage that genetic variants with a known functional role in creatinine production (or associated only with creatinine-based measurements of kidney function and not with cystatin-c or urea-based measurements) are unlikely to have true causal associations with kidney function decline over time. Associations with such variants in GWAS (ii) at the *GATM*, *CPS1*, and *SHROOM3* loci are likely to be due to bias and should be attenuated by an adjustment method which successfully reduces collider bias. They therefore can be used as a form of negative control. Conversely, associations with genetic variants at loci with probable true effects on kidney function decline over time (associated with multiple measures of kidney function or known implication of nearby genes in Mendelian kidney disease) should be relatively preserved.

We identified a set of independent genetic variants in the Gorski et al. summary statistics using the PLINK 1.9 “clump” algorithm^16^, using the p-value of cross-sectional eGFR associations from GWAS (i) as the clumping parameter with a clumping window of 100 kilobases and an *r^2^* threshold of 0.001.

The MR-Horse model was specified with the same structure and prior probability distributions as above (ie prior probability for _ placing most probability mass around a correlation between cross-sectional eGFR lowering effects, and faster eGFR decline), and posterior parameter estimates were drawn from four parallel Markov chains with 15,000 iterations and a burn in of 20%. In order to account for the fact that variants were selected for cross-sectional eGFR associations, and that the majority of such variants may have true associations with kidney function decline rendering a sparse horseshoe prior inappropriate, we performed a sensitivity analysis with less and more sparse prior probability distributions for (Supplemental data). Given that the true genetic architecture of many disease traits of interest is often unknown, sensitivity analyses were also performed using larger and smaller clumping windows for variant selection, and using deliberately mis-specified prior distributions for _ (Supplemental data).

## Results

### Simulations

Markov chains for β*_GY_* estimates converged satisfactorily (R −1.1 for > 99% of variants across 1000 iterations in each simulated scenario). Power for different adjustment methods by different simulated scenarios are shown in Figure 2a. The adapted MR-Horse model had less power to detect small variant effect sizes than unadjusted or Dudbridge method across the range of genetic correlations, but had greater power than the Slope-Hunter method when significant correlation was present. At the sample size tested, the power to detect variants of explaining ∼ 0.5% of the variance of the progression phenotype was 80-90% for the adapted MR Horse method across the range of genetic correlations. This range of effect size is relevant to kidney function phenotypes, with variants at the UMOD locus explaining ∼ 0.3 - 0.5% of the variance of eGFR decline^13^.

A collider bias effect was apparent in unadjusted analyses among variants with effects on disease incidence *G_I_* and *G_IP_* variants), and was strongest when genetic correlation was strongly positive (ie in the same direction as confounder effects +), and weakest when genetic correlation was negative (Fig 2b; unadjusted analyses). In the setting of no or little genetic correlation, this bias was reduced by the Dudbridge method, however the bias was markedly increased by both Dudbridge and Slope-Hunter methods in the setting of strong genetic correlation (Fig 2b). In our simulations the Slope-Hunter method increased the bias of *G_I_* and variants across the range of genetic correlations, likely due to ZeMRA violations. The MR13 Horse method reduced the bias of *G_I_* and *G_IP_* variants across the range of genetic correlations (Fig 2b).

The adapted MR-Horse consistently had lower Type 1 Error rates below the nominal (5%) level and below the other methods across the range of genetic correlations. This observation reflects the propensity of the method to shrink null effects to zero with a high precision whilst allowing non-zero effects to escape shrinkage.

Analyses with different sample sizes demonstrated a greater sensitivity of the power of the adapted MR-Horse method to sample size changes, with a larger proportional increase and drop-off in power with changes in sample size than the other methods (Supplemental figures 1-2).

Changes in the relative sizes of *G_I_*, *G_IP_* and *G_P_* clusters had significant effects on the performance of the Slope-Hunter and Dudbridge methods, with both methods acheiving comparable reductions in bias, and higher power than the adapted MR-Horse when the pleiotropic cluster was smaller (and explaining less phenotypic variance) than the non pleiotropic cluster. When cluster sizes were equal, the adapted MR-Horse achieved a superior reduction in bias and type 1 error in the setting of high genetic correlation (Supplemental figures 3-4). Relative performance of the adjustment methods was not significantly altered by changes in the parameters of the prior probability distributions for _ (Supplemental figures 5-6). Using a normal prior for *G_I_* effects resulted in greater power, but inflated bias and type 1 error rates, especially when correlation between *G_I_* and *G_IP_* was negative (Supplemental figure 7). The power of the MR-Horse method was greater when variants strongly associated with disease incidence also were strongly associated with disease progression (Supplemental Figure 8). In simulations we observed a positive relationship between power and F-statistics with a plateau at F statistics > 15 (referring to F statistics calculated based on observed disease progression associations among variants with significant disease incidence associations at p < 0.001).

### CKDGen data

Clumping yielded 531 independent genetic variants with available data in both GWAS (i) [GWAS of cross-sectional eGFR] and (ii) [GWAS of eGFR decline adjusted for baseline eGFR]. This pruning process removed ten of the lead variants from GWAS (ii) as they were not lead variants in GWAS (i); these variants were added back to the dataset giving a total set of 541 SNPs for use in the Dudbridge, Slope-Hunter and MR-Horse methods.

Application of the adapted MR-Horse method to summary statistics from the CKDGen GWAS of eGFR decline adjusted for baseline eGFR attenuated the magnitude of the estimates at all loci identified as significantly associated with eGFR decline (Fig 3). Credible intervals calculated by MR Horse were wider than confidence intervals in unadjusted estimates, reflecting the reduction in power associated with the method. Effect estimates for variants that were identified as likely to have a true effect on kidney function decline (notably at the *UMOD* locus) remained significantly associated with eGFR decline after MR-Horse adjustment (ie the 95% credible interval for did not include 0). For rs77924615 (considered to be a causal variant at the UMOD-PDILT locus^17^), the effect of one additional A allele on eGFR decline adjusted for baseline eGFR was 0.092 mls/min/year (95% confidence interval 0.081 - 0.103); adjusted using the adapted MR-Horse this was attenuated to 0.067 mls/min/year (equal-tailed 95% credible interval: 0.056 - 0.078). Associations at loci likely to have biased associations with eGFR decline (*GATM, CPS1* and *SHROOM3*) were not significantly associated with kidney function decline after MR-Horse adjustment. In the case of the *GATM* locus, where there is good evidence that the association is mediated directly by effects on creatine metabolism rather than kidney function^13^, the baseline-eGFR adjusted result was 0.029 mls/min/year per additional A allele (0.021 - 0.038); adjusted using the adapted MR-Horse model this was attenuated to 0.002 mls/min/year (−0.001 - 0.010). Sensitivity analyses with less and more sparse prior probabilities for β*_GY_*, alterations in clumping parameters or deliberately mis-specified prior distributions for did not alter the interpretation of estimates at GATM, CPS1 and SHROOM3 loci, although mis-specification of the probability priors did lead to likely true associations being rejected at the *ACVR2B, OVOL1* and *TPPP* loci (Supplemental Figures 9 - 11). Employing a normal prior for effects lead to marked shrinkage of estimates at loci with likely true effects (eg *UMOD*) whilst effects at loci likely to be biased (GATM, CPS1, SHROOM3) were not shrunk to zero (Supplemental figure 12).

**Figure 3:**
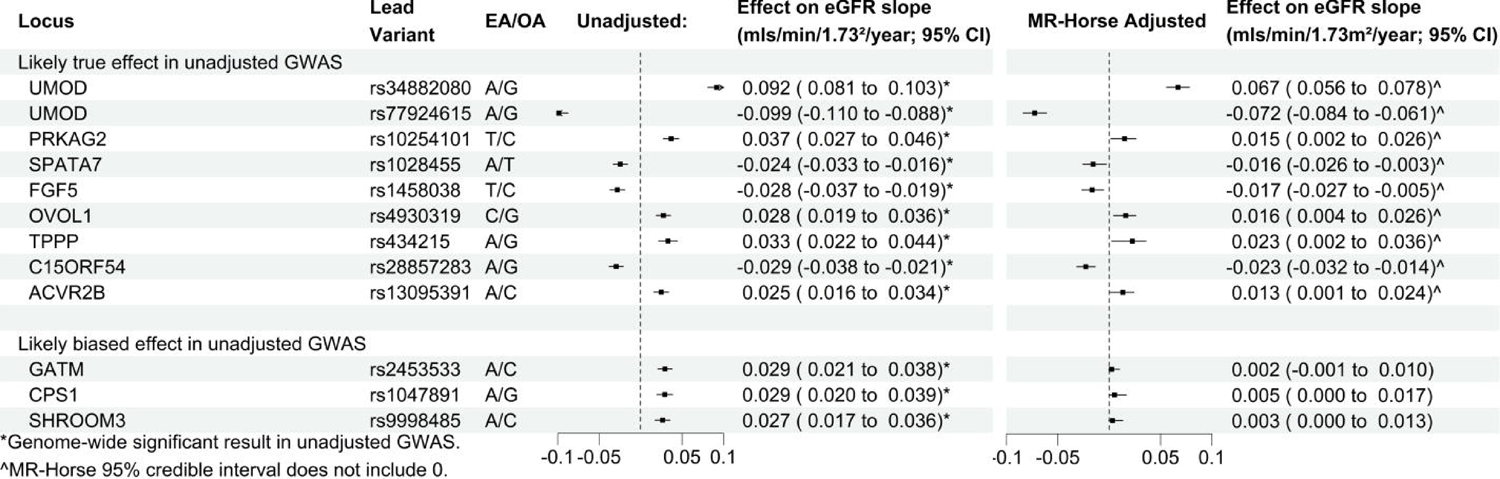
Unadjusted and MR-Horse adjusted associations of variants with eGFR decline, adjusted for baseline eGFR (adapted from CKDGen analyses). 1A: Associations of lead variants in Gorski et al.^13^ with eGFR decline, adjusted for baseline eGFR. Variants in top panel have likely true associations with eGFR-decline (associated with decline in analyses not adjusted for baseline eGFR). Variants in bottom panel have observed effects likely due to bias (not associated with eGFR decline in unadjusted analyses, and in the case of *GATM* and *CPS1*, likely related to creatinine metabolism directly rather than kidney function). 1B: Results after adjustment using adapted MR-Horse method. *Genome-wide significant result in GWAS adjusted for baseline eGFR. Significant result using MR-Horse method (95% credible interval does not include zero).

## Discussion

We set out to assess methods of studying the genetic associations of disease progression, accounting for effects of selection bias. The presented analyses have shown that an adapted version of the MR-Horse method appears to reliably reduce selection bias in GWAS of disease progression in simulated studies of phenotypes with heritability and polygenicity values within the range of values relevant to human complex traits^18^. When applied to real GWAS of kidney disease traits, it significantly attenuates associations at loci where currently observed associations are likely to be biased. Importantly, associations at loci with likely real effects are largely preserved, although attenuated in magnitude. This method may be useful in interpretation of signals in GWAS of disease progression where Type 1 error due to selection bias may be present.

The assumptions underlying the structure of the model used here are: (i) confounding is constant across genetic variants, (ii) genetic effects are independent and additive, (iii) all effects in Figure 1 are approximately linear and (iv) there are no direct effects of selected SNPs on confounders. Additionally, it is important that the definitions of disease incidence used in both GWAS of disease incidence and disease progression are as similar as possible. These assumptions are less restrictive than those of the Dudbridge or Slope-Hunter method, and notably there are no assumptions regarding the underlying correlation of CGX and CGY effects. assumptions are less restrictive than those of the Dudbridge or Slope-Hunter method, and This is important because the genetic architectures of disease progression traits are unknown a priori and methods that entail fewer assumptions are thus more useful for accurate inference.

Violations of assumption (i) could however arise if there was a single locus or small set of loci responsible for a large proportion of variance of either phenotype. This would mean that the genetic component of *U* would vary depending on whether a variant being tested was at or in responsible for a large proportion of variance o either phenotype. This would mean that the LD with such a locus. This could be ameliorated by removing such a locus from the data, and could still be estimated from such an analysis. Violation of assumption (ii) is possible, but non-additive effects are likely a small contributor to heritability of complex traits^19^. Assumption (iii) is likely to hold when the effects concerned are small. Assumption (iv) can be tested for known confounders with existing GWAS evidence.

Results of this method will be to some extent influenced by the prior probability distributions prior) for the C_GYi_ effects in our analysis is inappropriate if we consider pleiotropy to be a major specified by the user. It could be argued that using a horseshoe prior (or alternative shrinkage characteristic of the genetic architecture of disease progression, and some work has shown that posterior estimates can be very sensitive to hyperprior choice when using horseshoe priors^20^.

We have shown in sensitivity analyses that use of a normal prior for C_GY_ effects resulted in inflated type 1 error rates in simulations, and a failure to reject the likely biased associations at *CPS1, GATM* and *SHROOM3* in CKDGen data. Importantly, type 1 error rates when using a normal prior were sensitive to the magnitude of correlation between genetic effects on incident and progressive disease; this is an undesirable characteristic when the true underlying correlation is unknown. Results of our simulations and the CKDGen analyses were not significantly changed despite large magnitude alterations of (i) the sparseness of the horseshoe prior distribution and (ii) the prior probability for the correlation of genetic effects. Using the horseshoe prior therefore retains sufficient power to detect important effects in real GWAS, whilst biased effects are shrunk to zero.

In practice, it will computationally intractable for most users to fit the MR-Horse model on a genome wide scale. As such, a subset of variants must be selected on which the model should be fitted. Specific considerations when selecting these variants are (a) they must be independent, (b) they should be strongly associated with disease incidence and (c) they should explain a sizeable proportion of the variance of disease progression. We suggest based on simulations that an F statistic > 15 calculated with respect to the observed disease progression associations should provide adequate power for the MR-Horse method.

A further limitation of the method is the fact that the bias reductions are small relative to unadjusted analyses in the setting of no or negative genetic correlation (or in the more general case, where the correlation of genetic effects is in the opposite direction to the correlation of confounder effects responsible for the collider bias). The power reduction when using MR-Horse in these analyses is however quite marked. It may be preferable therefore in situations where correlation appears unlikely to use unadjusted estimates or alternative methods for adjustment. For example in the case of cancer where incident disease is driven by cell proliferation, but disease progression driven by invasion and metastasis, it is possible that the biology and genetic architecture of disease progression and incidence are distinct^21^.

Assumptions of no or low correlation may therefore be appropriate. However we again emphasise that in most cases the true underlying correlation structure is unknown, and the relatively consistent performance of MR-Horse across the range of correlations renders it less vulnerable to model mis-specification. Unadjusted analyses and Slope-Hunter and Dudbridge methods can produce extreme biases when significant correlation is present.

In principal, this method could be applied to binary outcomes (for example incident end-stage kidney disease among those with a specific kidney disease type), although similar to instrumental variable analyses such as MR, such estimates may become biased towards the null method to observational epidemiology more broadly. The coefficient in Figure 1D, which due to the non-collapsibility of odds ratios^22^. There are also potential applications of this defines the magnitude of any collider bias effect and can be estimated from the MR-Horse model, is independent of the specific exposure of interest and could be applied to reduce bias of estimated effects on disease progression in other observational studies provided that “incident disease” and “progression” are defined in the same manner as in the genetic study of interest.

Unbiased estimates of the associations of genetic and environmental factors with disease prognosis are crucial to understanding disease mechanisms and predicting disease risks. As drug development pipelines become more reliant on genetic evidence for target selection^23^, it is increasingly important that the genetic evidence base is unbiased. This adaptation of the MR-Horse method minimises selection bias, which is an under-addressed source of error in studies of disease progression.

## Data availability

Raw simulated datasets used in this work are available from the authors on reasonable request. CKDGen summary statistics are publicly available for download from the CKDGen website.

## Code availability

Code will published in a public GitHub repository on publication of the manuscript

## Author contributions

KD, NS and WGH conceived and designed the study. KD, JT and NS conducted the statistical analyses. KD, JT, DZ, WGH and NS drafted and revised the manuscript.

## Ethical approval

No ethical approval was required for this work as we used only simulated data or publicly available summary statistics.

## References

1 Galesloot TE, Grotenhuis AJ, Kolev D, Aben KK, Bryan RT, Catto JWF et al. Genome-wide Meta-analysis Identifies Novel Genes Associated with Recurrence and Progression in Non-muscle-invasive Bladder Cancer. European Urology Oncology 2022; 5: 70–83.

2 Fogh I, Lin K, Tiloca C, Rooney J, Gellera C, Diekstra FP et al. Association of a Locus in the CAMTA1 Gene With Survival in Patients With Sporadic Amyotrophic Lateral Sclerosis. JAMA neurology 2016; 73: 812–820.

3 Chang I-S, Jiang SS, Yang JC-H, Su W-C, Chien L-H, Hsiao C-F et al. Genetic Modifiers of Progression-Free Survival in Never-Smoking Lung Adenocarcinoma Patients Treated with First-Line Tyrosine Kinase Inhibitors. American Journal of Respiratory and Critical Care Medicine 2017; 195: 663–673.

4 Hernán MA, Hernández-Díaz S, Robins JM. A Structural Approach to Selection Bias. Epidemiology 2004; 15: 615.

5 Dahabreh IJ, Kent DM. Index event bias: An explanation for the paradoxes of recurrence risk research. JAMA: The Journal of the American Medical Association 2011; 305: 822–823.

6 Mitchell RE, Hartley AE, Walker VM, Gkatzionis A, Yarmolinsky J, Bell JA et al. Strategies to investigate and mitigate collider bias in genetic and Mendelian randomisation studies of disease progression. PLoS genetics 2023; 19: e1010596.

7 Dudbridge F, Allen RJ, Sheehan NA, Schmidt AF, Lee JC, Jenkins RG et al. Adjustment for index event bias in genome-wide association studies of subsequent events. Nature Communications 2019; 10: 1561.

8 Cai S, Hartley A, Mahmoud O, Tilling K, Dudbridge F. Adjusting for collider bias in genetic association studies using instrumental variable methods. Genetic Epidemiology 2022; 46: 303– 316.

9 Robinson-Cohen C, Triozzi JL, Rowan B, He J, Chen HC, Zheng NS et al. Genome-Wide Association Study of CKD Progression. Journal of the American Society of Nephrology: JASN 2023; 34: 1547–1559.

10 Mahmoud O, Dudbridge F, Davey Smith G, Munafo M, Tilling K. A robust method for collider bias correction in conditional genome-wide association studies. Nature Communications 2022; 13: 619.

11 Grant AJ, Burgess S. A Bayesian approach to Mendelian randomization using summary statistics in the univariable and multivariable settings with correlated pleiotropy. The American Journal of Human Genetics 2024; 111: 165–180.

12 Carvalho CM, Polson NG, Scott JG. The horseshoe estimator for sparse signals. Biometrika 2010; 97: 465–480.

13 Gorski M, Rasheed H, Teumer A, Thomas LF, Graham SE, Sveinbjornsson G et al. Genetic loci and prioritization of genes for kidney function decline derived from a meta-analysis of 62 longitudinal genome-wide association studies. Kidney International 2022; 102: 624–639.

14 Hardy GH. MENDELIAN PROPORTIONS IN A MIXED POPULATION. *Science (New York*, NY) 1908; 28: 49–50.

15 Wuttke M, Li Y, Li M, Sieber KB, Feitosa MF, Gorski M et al. A catalog of genetic loci associated with kidney function from analyses of a million individuals. Nature Genetics 2019; 51: 957–972.

16 Purcell S, Neale B, Todd-Brown K, Thomas L, Ferreira MAR, Bender D et al. PLINK: A Tool Set for Whole-Genome Association and Population-Based Linkage Analyses. The American Journal of Human Genetics 2007; 81: 559–575.

17 Gorski M, Jung B, Li Y, Matias-Garcia PR, Wuttke M, Coassin S et al. Meta-analysis uncovers genome-wide significant variants for rapid kidney function decline. Kidney international 2021; 99: 926–939.

18 Zeng J, de Vlaming R, Wu Y, Robinson Matthew R et al. Signatures of negative selection in the genetic architecture of human complex traits. Nature Genetics 2018; 50: 746–753

19 Zhu Z, Bakshi A, Vinkhuyzen Anna AE, Hemani G, Lee S, Nolte Ilja M et al. Dominance genetic variation contributes little to the missing heritability for human complex traits. American Journal of Human Genetics 2015; 96: 377–385.

20 Piironen J, Vehtari A. On the hyperprior choice for the global shrinkage parameter in the horseshoe prior. Artificial Intelligence and Statistics 2017;: 905–913.

21 Chathrath A, Ratan A, Dutta A. Germline variants that affect tumor progression. Trends in Genetics 2021; 37(5) 433 – 443

22 Burgess S, Small D, Thompson S. A review of instrumental variable estimators for Mendelian Randomization. Statistical Methods in Medical Research 2015; 26(5)

23 Minikel EV, Painter JL, Dong CC, Nelson MR. Refining the impact of genetic evidence on clinical success. Nature 2024; 629: 624–629.

