## Supplementary material for "An application of the MR-Horse method to reduce selection bias in genome-wide association studies of disease progression": Data supplement

K Donovan et al.

Data supplement

Section 1: Supplemental simulations	Page 2

Section 2: CKDGen Sensitivity analyses	Page 10

**Section 1: Supplemental simulation analyses**

Supplemental Figure 1: Results from simulations with sample size 5,000

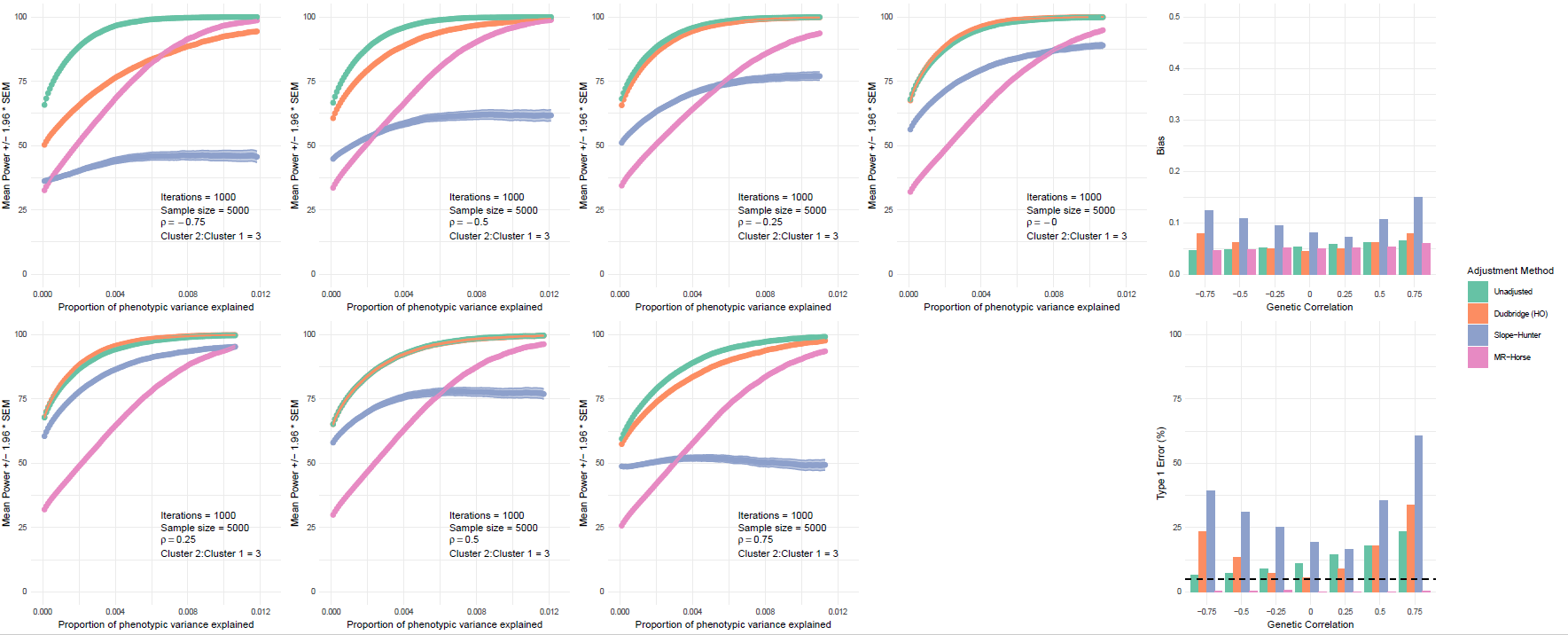

Left: Power of unadjusted and adjusted analyses to detect $\beta_{GY}$ effects across the range of effect sizes, by different values of $\rho$. Plotted values are mean power (%) across 1,000 simulation iterations with shaded areas +/- 1.96 * SEM. Phenotypic variance explained defined as: $2\bar{\beta}^{2}maf\left( 1-maf \right)$. Right: Bias (top panel) and type 1 error rates (bottom panel) of each method (mean bias of $G_{I}$ and $G_{IP}$ variants across 1,000 simulation iterations).

Supplemental Figure 2: Results from simulations with sample size 20,000

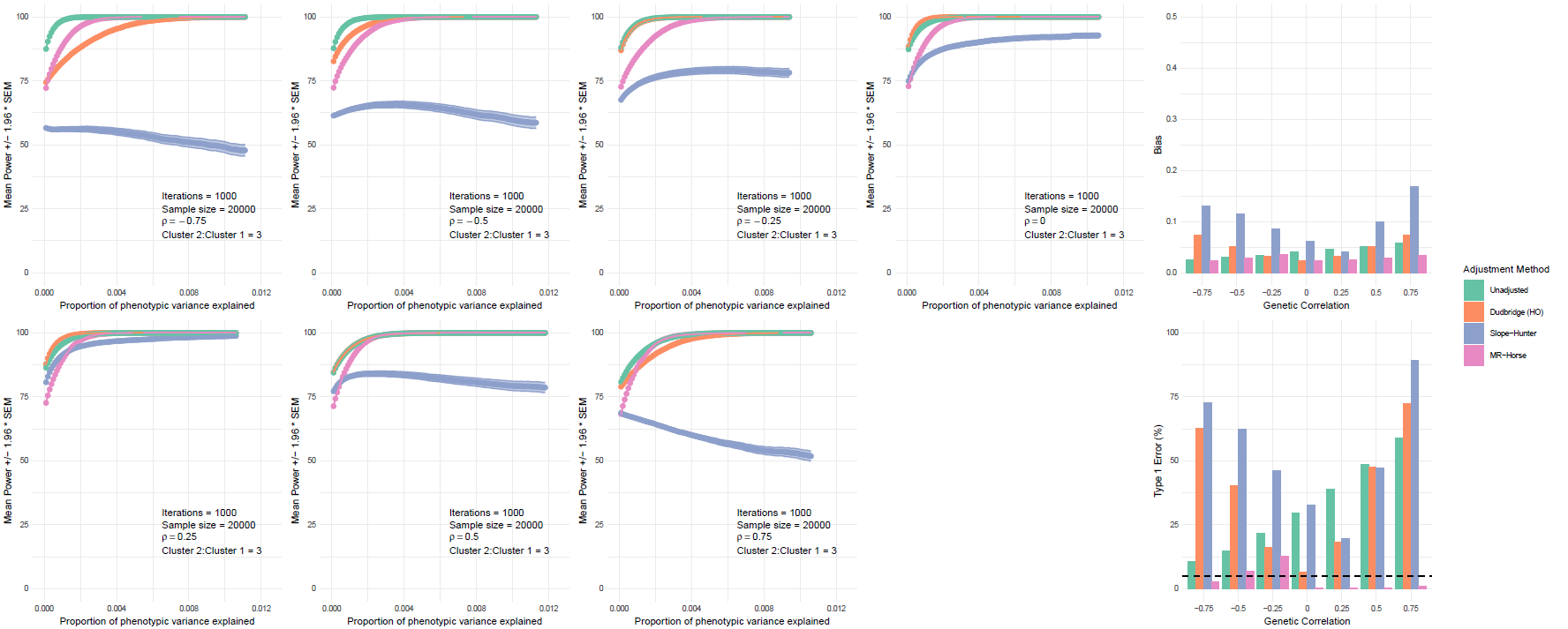

Left: Power of unadjusted and adjusted analyses to detect $\beta_{GY}$ effects across the range of effect sizes, by different values of $\rho$. Plotted values are mean power (%) across 1,000 simulation iterations with shaded areas +/- 1.96 * SEM. Phenotypic variance explained defined as: $2\bar{\beta}^{2}maf\left( 1-maf \right)$. Right: Bias (top panel) and type 1 error rates (bottom panel) of each method (mean bias of $G_{I}$ and $G_{IP}$ variants across 1,000 simulation iterations).

Supplemental Figure 3: Results from simulations with equal sized $G_{I}$ and $G_{IP}$ clusters

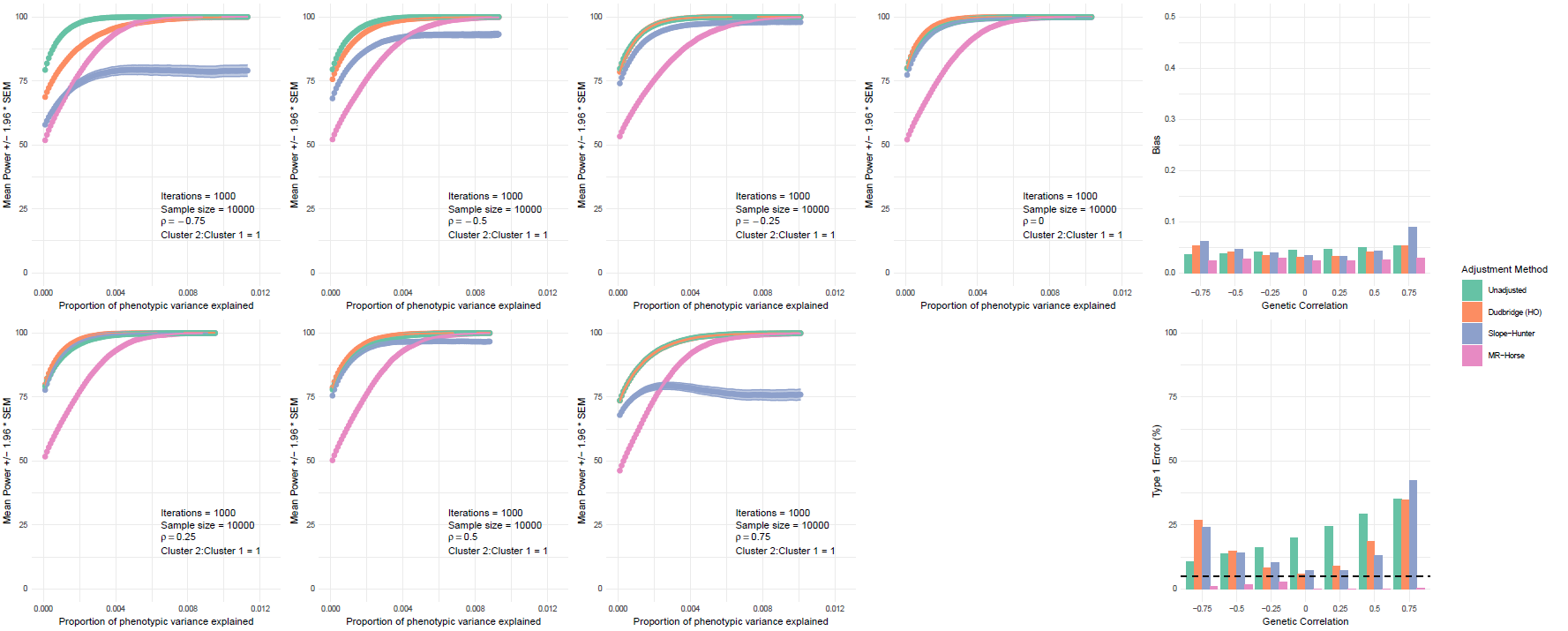

Left: Power of unadjusted and adjusted analyses to detect $\beta_{GY}$ effects across the range of effect sizes, by different values of $\rho$. Plotted values are mean power (%) across 1,000 simulation iterations with shaded areas +/- 1.96 * SEM. Phenotypic variance explained defined as: $2\bar{\beta}^{2}maf\left( 1-maf \right)$. Right: Bias (top panel) and type 1 error rates (bottom panel) of each method (mean bias of $G_{I}$ and $G_{IP}$ variants across 1,000 simulation iterations). $G_{I}$ and $G_{IP}$ clusters equal in size and each explaining 20% of the phenotypic variance.

Supplemental figure 4: Results from simulations with larger $G_{I}$ than $G_{IP}$ cluster

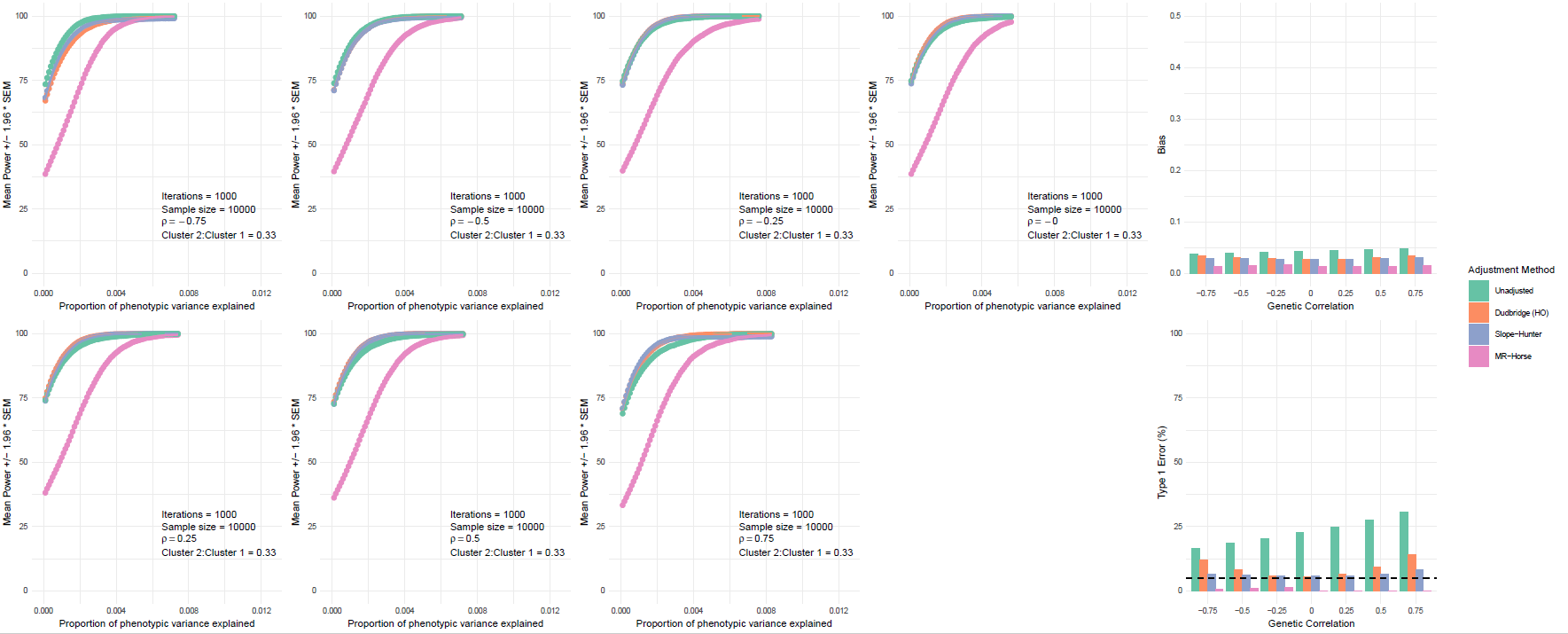

Left: Power of unadjusted and adjusted analyses to detect $\beta_{GY}$ effects across the range of effect sizes, by different values of $\rho$. Plotted values are mean power (%) across 1,000 simulation iterations with shaded areas +/- 1.96 * SEM. Phenotypic variance explained defined as: $2\bar{\beta}^{2}maf\left( 1-maf \right)$. Right: Bias (top panel) and type 1 error rates (bottom panel) of each method (mean bias of $G_{I}$ and $G_{IP}$ variants across 1,000 simulation iterations). $G_{I}$ cluster 3 times larger than $G_{IP}$ cluster. Clusters explain 30% and 10% respectively of the phenotypic variance.

Supplemental Figure 5: Results from simulations with less sparse prior for $\beta_{GY}$

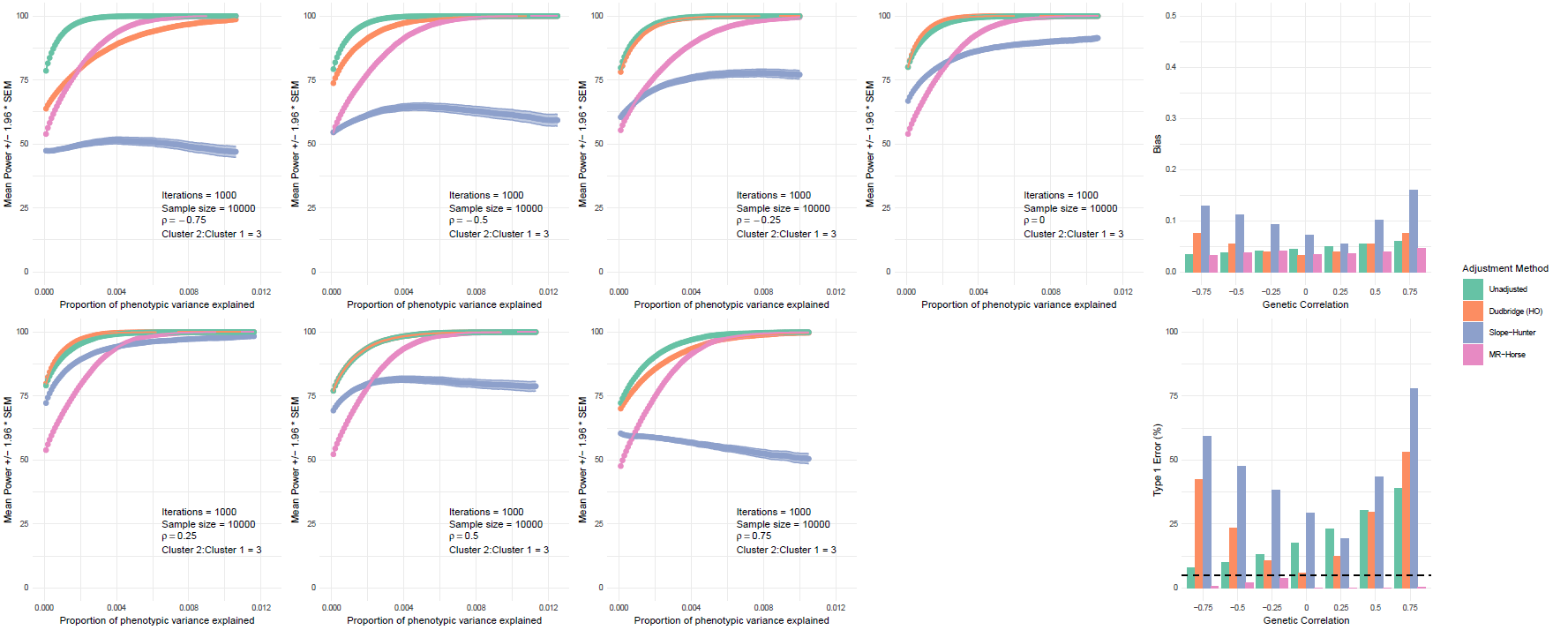

Left: Power of unadjusted and adjusted analyses to detect $\beta_{GY}$ effects across the range of effect sizes, by different values of $\rho$. Plotted values are mean power (%) across 1,000 simulation iterations with shaded areas +/- 1.96 * SEM. Phenotypic variance explained defined as: $2\bar{\beta}^{2}maf\left( 1-maf \right)$. Right: Bias (top panel) and type 1 error rates (bottom panel) of each method (mean bias of $G_{I}$ and $G_{IP}$ variants across 1,000 simulation iterations). Prior probability distribution for $\beta_{GY}$ less sparse than in main analysis – this is achieved by specifying a prior distribution for the global shrinkage parameter tau of Cauchy^+^(0,100).

Supplemental Figure 6: Results from simulations with more sparse prior for $\beta_{GY}$

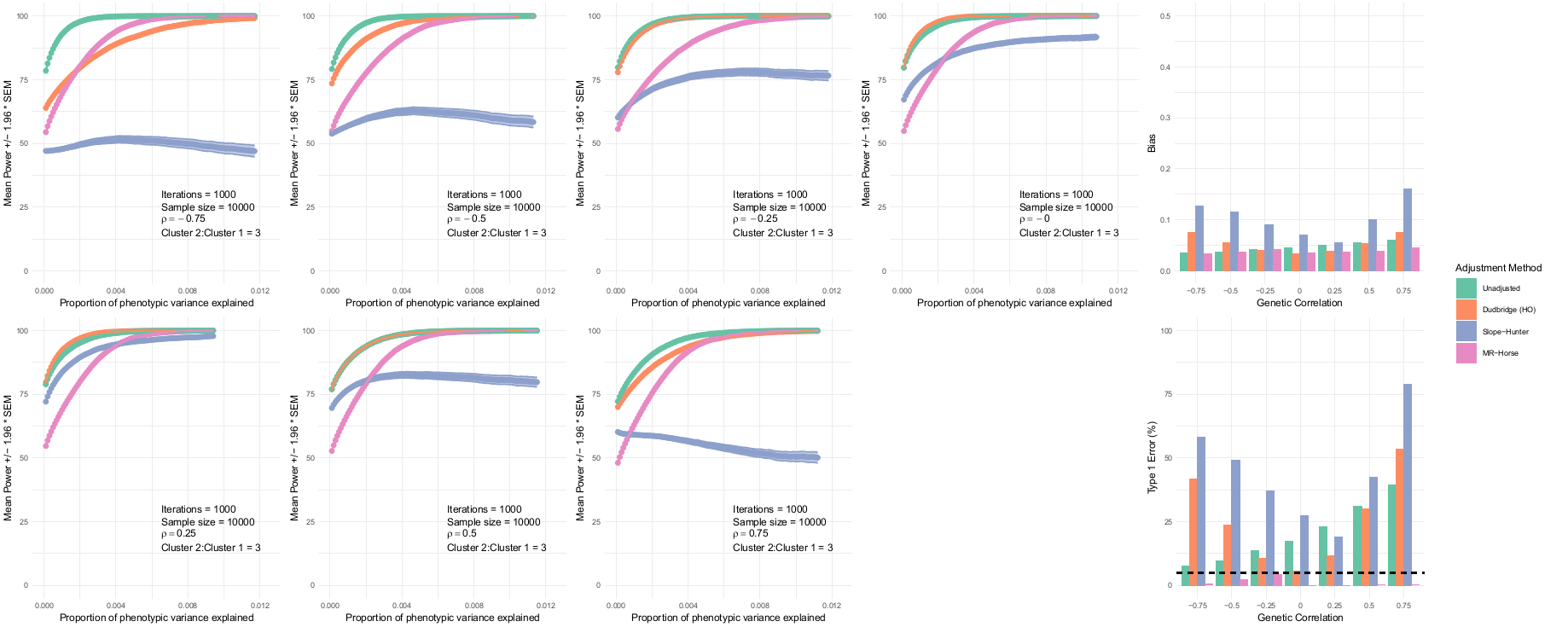

Left: Power of unadjusted and adjusted analyses to detect $\beta_{GY}$ effects across the range of effect sizes, by different values of $\rho$. Plotted values are mean power (%) across 1,000 simulation iterations with shaded areas +/- 1.96 * SEM. Phenotypic variance explained defined as: $2\bar{\beta}^{2}maf\left( 1-maf \right)$. Right: Bias (top panel) and type 1 error rates (bottom panel) of each method (mean bias of $G_{I}$ and $G_{IP}$ variants across 1,000 simulation iterations). Prior probability distribution for $\beta_{GY}$ more sparse than in main analysis – this is achieved by specifying a prior distribution for the global shrinkage parameter tau of Cauchy^+^(0,0.01).

Supplemental Figure 7: Results from simulations with a normal prior for $\beta_{GY}$

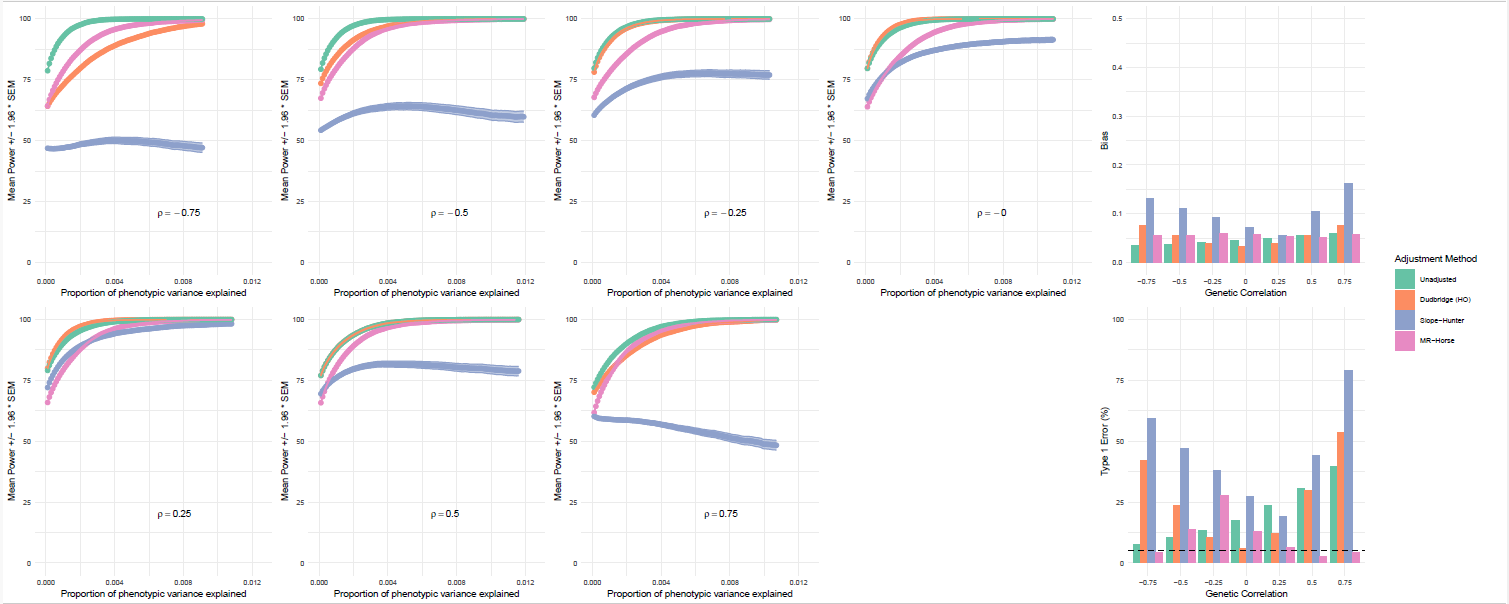

Left: Power of unadjusted and adjusted analyses to detect $\beta_{GY}$ effects across the range of effect sizes, by different values of $\rho$. Plotted values are mean power (%) across 1,000 simulation iterations with shaded areas +/- 1.96 * SEM. Phenotypic variance explained defined as: $2\bar{\beta}^{2}maf\left( 1-maf \right)$. Right: Bias (top panel) and type 1 error rates (bottom panel) of each method (mean bias of $G_{I}$ and $G_{IP}$ variants across 1,000 simulation iterations). Prior probability distribution for $\beta_{GY}$ follows a normal distribution with mean zero and a non-informative prior for the variance.

Supplemental Figure 8: Power of MR-Horse method is positively associated with F-statistic of variant-progression associations

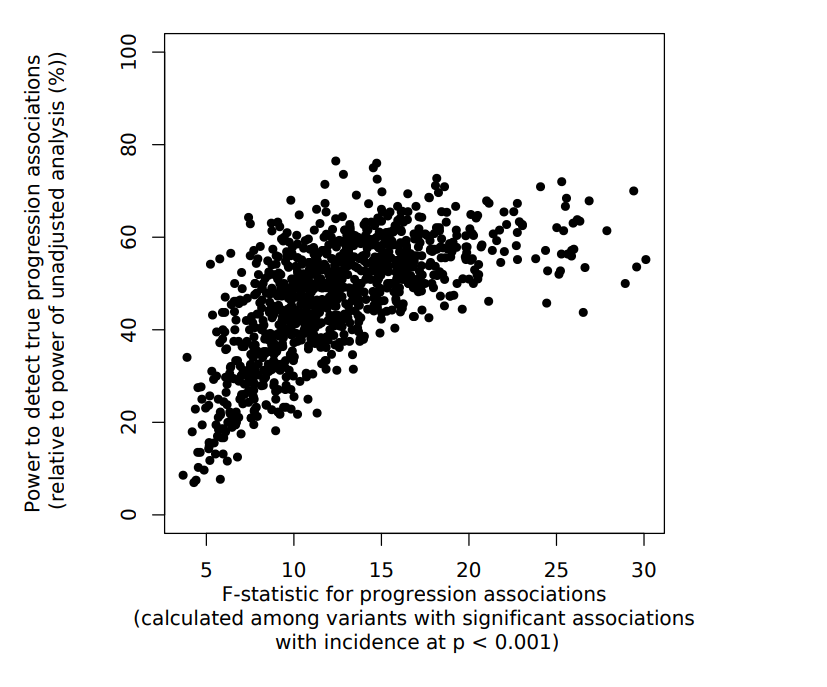

**Section 2: CKDGen Sensitivity analyses**

Supplemental Figure 9: Results from CKDGen analyses under mis-specification of prior probability distribution for effect correlation

A: Uniform prior in the range -1 – 1

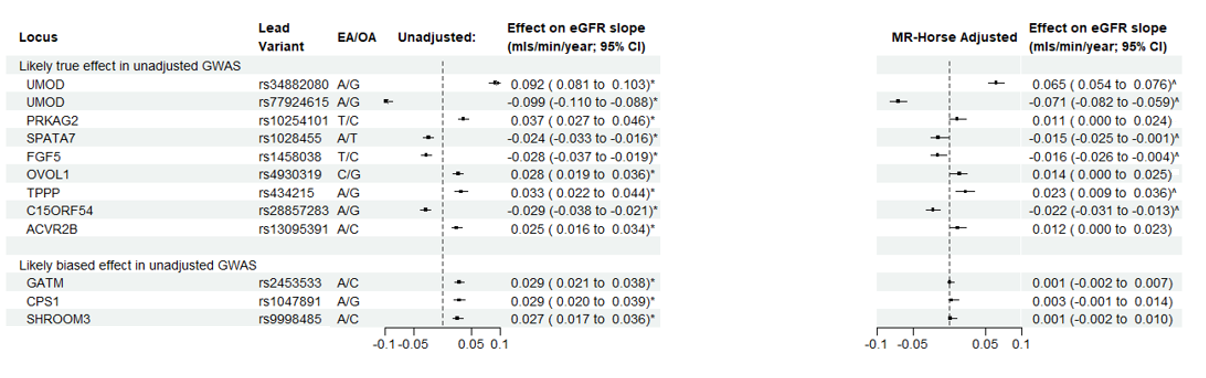

B: Inversely specified prior, with most mass around a probability of correlation between eGFR raising alleles and faster eGFR decline
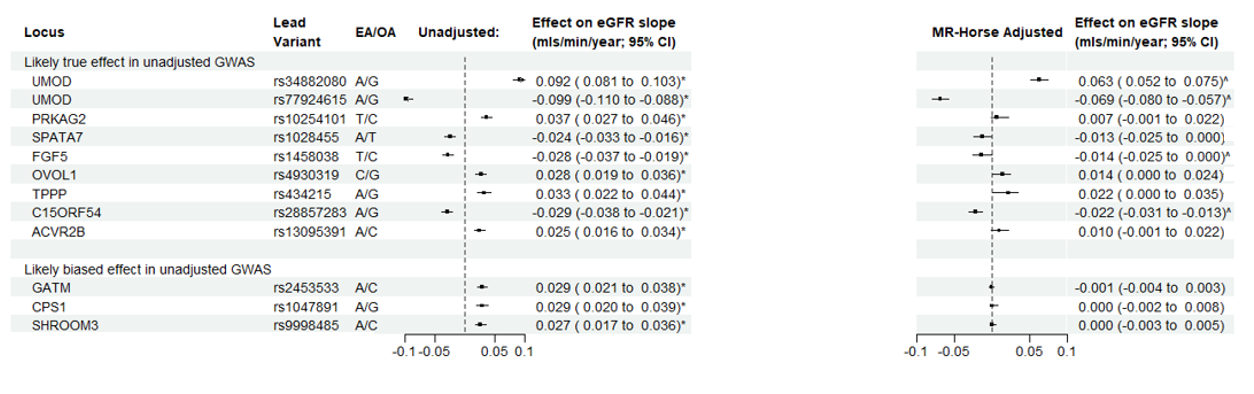

*Genome-wide significant result in GWAS adjusted for baseline eGFR. ^Significant result using MR-Horse method (95% credible interval does not include zero).

Supplemental Figure 10: Results from CKDGen analyses under sparser and less sparse prior for $\beta_{GY}$

A: Less sparse prior probability for $\beta_{GY}$ (achieved by specifying prior for global shrinkage parameter of Cauchy^+^(0, 100))

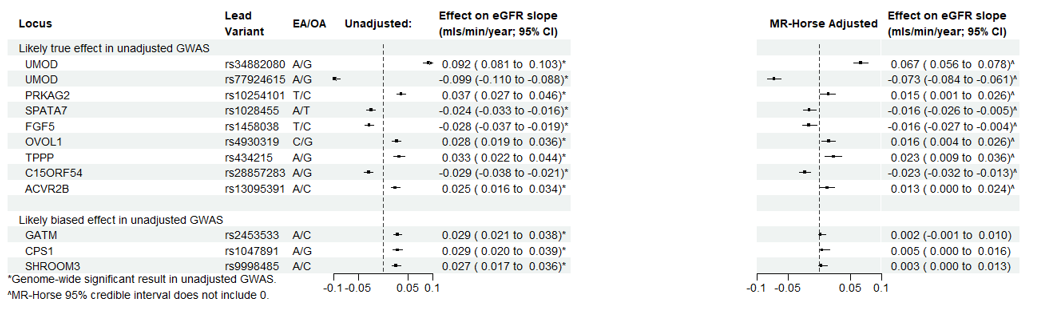

B: More sparse prior probability for $\beta_{GY}$ (achieved by specifying prior for global shrinkage parameter of Cauchy^+^(0, 0.01))

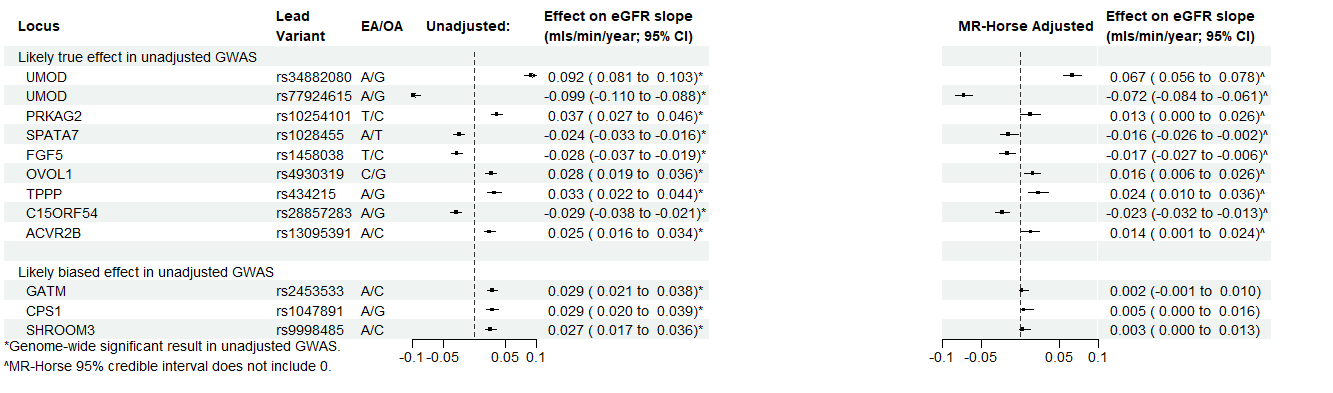

Supplemental Figure 11: Results from CKDGen analyses under with different clumping parameters used for variant selection

A: Narrow clumping window used (1kb)

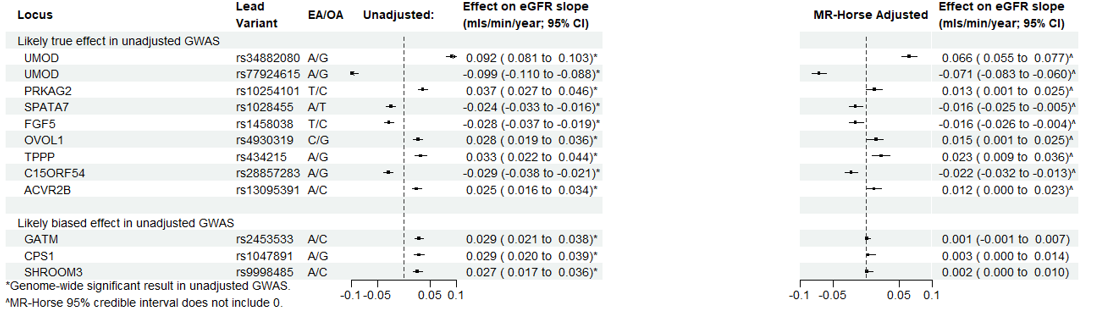

B: Wide clumping window used (10Mb)

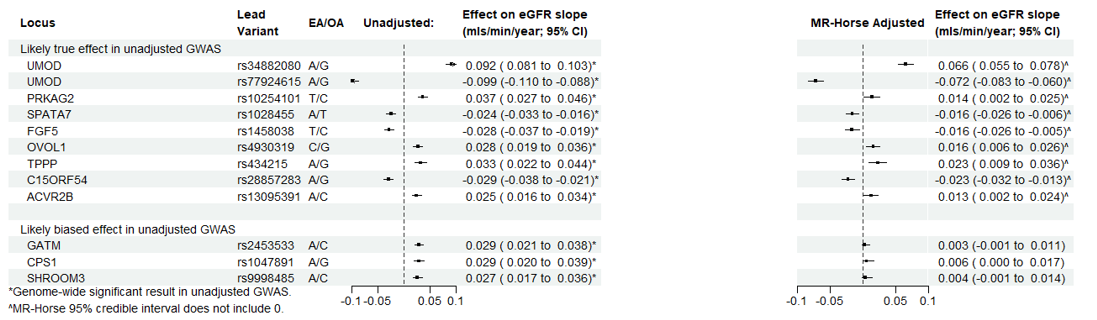

Supplemental Figure 12: Results from CKDGen analyses under a normal prior for $\beta_{GY}$

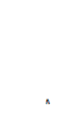

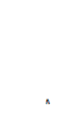

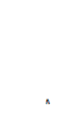

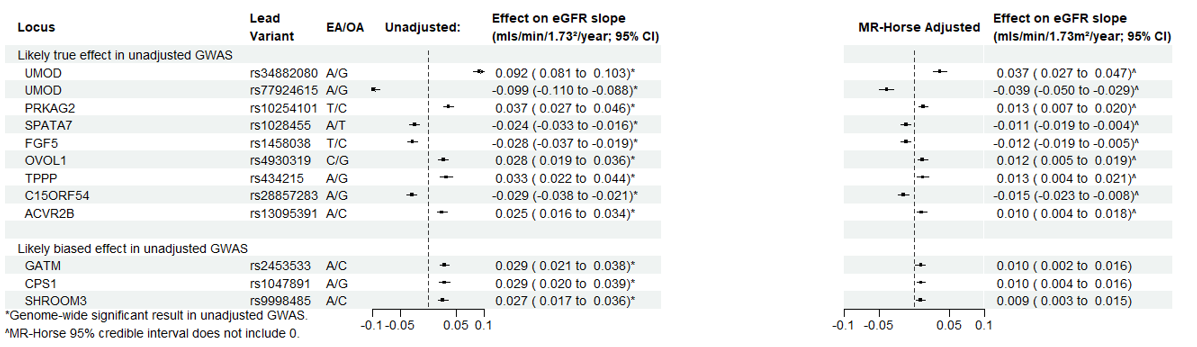
